# Kinetic analysis methods applied to single motor protein trajectories

**DOI:** 10.1101/287748

**Authors:** A L Nord, A F Pols, M Depken, F Pedaci

## Abstract

Molecular motors convert chemical or electrical energy into mechanical displacement, either linear or rotary. Under ideal circumstances, single-molecule measurements can spatially and temporally resolve individual steps of the motor, revealing important properties of the underlying mechanochemical process. Unfortunately, steps are often hard to resolve, as they are masked by thermal noise. In such cases, details of the mechanochemistry can nonetheless be recovered by analyzing the fluctuations in the recorded traces. Here, we expand upon existing statistical analysis methods, providing two new avenues to extract the motor step size, the effective number of rate-limiting chemical states per translocation step, and the compliance of the link between the motor position and the probe particle. We first demonstrate the power and limitations of these methods using simulated molecular motor trajectories, and we then apply these methods to experimental data of kinesin, the bacterial flagellar motor, and F_1_-ATPase.

## 1 Intro

Molecular motors convert chemical and electrical energy into mechanical work in the form of linear or rotary motion. This process can be seen as a stochastic sequence of chemical and conformational changes driven by chemical-energy consumption and influenced by thermal forces. While bulk measurements are able to report on the average of the stochastic motion, they fail to capture the information contained in the single-molecule fluctuations. However, advances in single molecule techniques, which provide the opportunity to measure the movement of single motor proteins *in vivo* and *in vitro*, have proven remarkably valuable in the study of molecular motors. Given adequate spatio-temporal resolution, the mechanochemical reaction schemes of motor proteins can be resolved, providing details of the underlying microscopic mechanism [1]. Single molecule experiments have enabled the direct resolution of individual steps in many molecular motors, revealing details of the mechanochemical cycle for kinesin [2,3], myosin [4–7], dynein [8,9], F_1_-ATPase [10, 11], the viral packaging motor *ϕ*29 [12, 13], and the bacterial flageller motor [14]. Such measurements yield information about the fundamental mechanical step size, the number of chemical processes per mechanical step, the transition rate between states, and the number of fuel molecules consumed per mechanochemical cycle.

Despite great experimental advances, the signal-to-noise ratio of single motor trajectories is often insufficient to resolve individual motor steps. Under these conditions, analysis of the stochastic fluctuations can provide information not deducible from average quantities. For example, in the time domain, measurements of the variance in displacement have yielded the number of rate-limiting processes and the number of ATP molecules consumed per mechanochemical cycle of kinesin [15–17]. In the frequency domain, the underlying stochastic signal and the noise often have separable power spectra, and can thus be identified and characterized in Fourier space, yielding the number of rate-limiting steps in the enzymatic cycle [18]. Such analysis has further yielded the step size of UvrD helicase [19] and DNA translocase FtsK [20], and the relaxation step of DNA supercoiling type II topoisomerase [18].

In this study, we extend previous work in order to provide analytical solutions to i) the steady-state variance of a reporter-particle displacements over fixed time intervals and ii) the power spectrum of the speed of the reporter particle. Under certain conditions, these methods, one of which operates in the time domain and the other in the frequency domain, are capable of recovering the mechanical step size of the motor, the number of chemical processes per step, and the stiffness of the linker between the motor protein and probe, even under conditions where the individual steps are not spatially and temporally resolved. We perform simulations to compare the performance of these methods, and we demonstrate their utility in extracting additional information from previously published experimental data of three molecular motors.

## 2 Theoretical analysis of single motor trajectories

In order to extract additional information from the stochastic fluctuations of a reporter particle, we need to model the effects of the underlying mechanochemical cycle on its motion. In a classical approach to macromolecular dynamics [15], which shares several aspects with the treatment of shot noise in physical systems [21, 22], the displacement *x*_b_(*t*) of the observable reporter object, often a microscopic bead, tethered to the molecular motor is modeled by

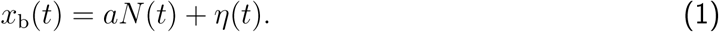

Here *a* is the size of the mechanical step of the motor along its track, *N* (*t*) is the total number of mechanical steps in time *t*, and *η* is the thermal noise characterized by 〈*η*〉 = 0 and 〈*η*^2^〉 = *k*_B_*T/k*_*l*_, where *k*_*l*_ is the total stiffness of the construct. Implicit in this description is further that the response of the system is faster than any other relevant timescale of the system, including the motor stepping rate. This condition might not always be clearly satisfied *a priori*, and we relax this condition by instead modeling the bead motion as governed by Stoke’s drag (drag coefficient *γ*)

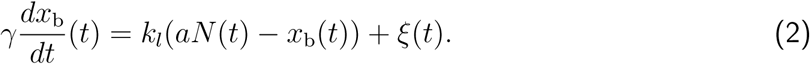

where the random force *ξ*(*t*) satisfies [23]

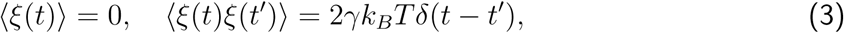

and *δ* is the Dirac-delta function.

### 2.1 Time-domain analysis of reporter-particle position fluctuations

In the case that the motor is a Poisson stepper, and the reporter particle instantaneously responds to motor displacements (i.e. when Equation (1) holds), both the average and variance of the reporter-particle displacement increases linearly with time in the steady state [15]

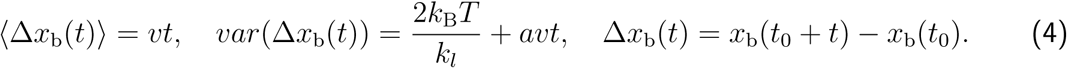

The utility of this approach comes from the fact that Eq (4) can be used to estimate the step size *a* and construct stiffness *k*_*l*_ by analyzing the average and variance of the experimentally accessible reporter-particle displacements. More precisely, Eq (4) shows that

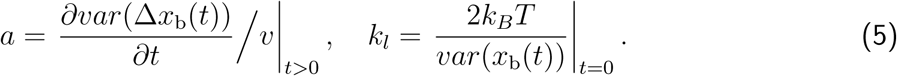

More generally, if the step size is known, the randomness parameter

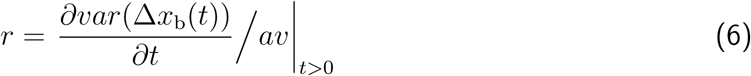

has been introduced [15] to quantify how much the process deviates from Poisson statistics (*r* = 1). In the simple case where only every m:th chemical step gives rise to translocation, it can be shown that *r* = 1/*m*, and if we know the step size we can estimate how many effective chemical steps there is in one translocation cycle [15].

As we aim to also capture the relaxation dynamics of the system, we extend the above analysis by considering the variance and average displacement of a reporter particle that moves according to Eq. (2). Assuming that the molecular motor is a Poisson stepper, wherein each mechanical step of the motor is a consequence of a single rate-limiting biochemical process we find that the variance of the bead position can be expressed as (see derivation in the SI)

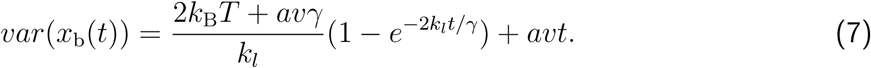

It is worth noting that this expression of the variance expands the classical result described above by Eq (4), including the filtering effect of the elastic linkage, which imposes a characteristic relaxation time *t*_*c*_ = *γ/k*_*l*_. In the limit of *t*_*c*_ → 0, Eq (4) is retrieved from Eq (7). Therefore, by measuring the bead position *x*_b_(*t*), its variance *var*(*x*_b_(*t*)) can be calculated and fit with Eq (7) to obtain the relaxation time (or equivalently the drag), as well as the step size *a* and the construct stiffness *k*_*l*_. Fig S1F-I demonstrate the behavior of Eq (7) as a function of motor step size, motor speed, stiffness of the linker, and bead size. It is interesting to note that if the average energy dissipated by moving the bead a distance equal to the motor step (*avγ*) is not negligible compared to twice the thermal energy (2*k*_B_*T*), then we will underestimate the construct stiffness if we assume the dynamics to be governed by Eq. (1).

### 2.2 Frequency-domain analysis of reporter-particle velocity fluctuations

Instead of analyzing the motor position in the time domain, we now approach the molecular system by analyzing the speed of the motor in the frequency domain. In the time domain, characteristic behaviors on different timescales are convoluted, and often hard to disentangle. In the frequency domain such time-separated behaviors are often readily disentangled as they affect different frequency ranges. This is true, for instance, for low signal-to-noise measurements where the signal and noise, due to their statistics, can by separated and identified in the Fourier domain [18]. Below we show that an analysis of the speed of the motor in the frequency domain provides the interesting ability of simultaneously quantifying step size, construct stiffness, and relaxation time of the molecular system. In the frequency domain we further manage to extend our considerations to the situation where only every m:th chemical cycle results in translocation, allowing us to estimate also the effective number of rate-limiting chemical steps for each translocation step.

Taking the Fourier transform of Eq (2) (derived in SI), the power spectral density (PSD) of the speed of the bead velocity (*v*_b_ = *dx*_b_/*dt*) as a function of frequency *f* (measured in Hz) can be written as

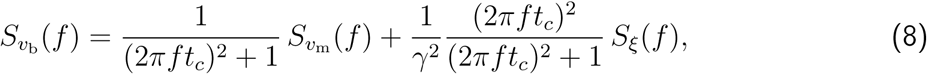

where *t*_*c*_ = *γ/k*_*l*_ is the characteristic relaxation time of the construct. The two components 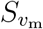 and *S*_*ξ*_ are the PSD of the speed of the motor *v*_m_, and the PSD of the thermal forces *ξ* acting on the bead, respectively. By direct analogy to shot noise in electronic circuits (see SI), the PSD of the motor’s velocity can be written as

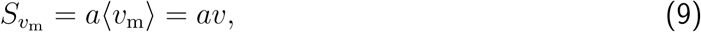

in the steady state. We note here that 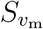 is independent of frequency. The PSD of the thermal forces acting on the bead can be computed by applying the Wiener-Khinchin theorem [24] to Eqs (3), giving

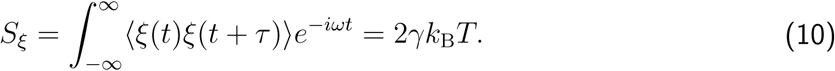

Note that the above term is independent of frequency as well, i.e. the thermal forces acting on the bead produce white noise. Finally, combining Eqs (8), (9), and (10), the explicit analytical expression for the PSD of the speed of a bead *v*_*b*_ elastically tethered to a Poissonian stepping motor can be written as

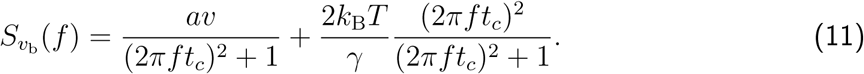

This result can be generalized to the situation where only every m:th chemical step gives rise to translocation of the motor. Letting *τ* be the average time between translocation events, we find that the PSD of the motor speed, gains an extra factor *α* (derived in the SI), 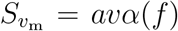, with

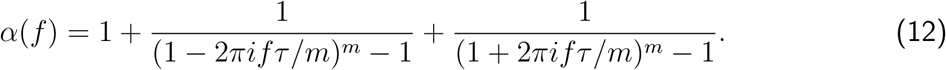

Therefore in this more general case, the overall PSD of the speed of the bead becomes

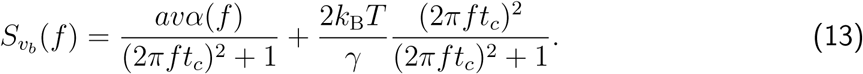

We note that in the case of a pure Poisson stepper (*m* = 1), *α* = 1 and Eq (13) reduces to Eq (11). Fig S1A-E demonstrate the behavior of Eq (13) as a function of motor step size, motor speed, construct stiffness, probe size, and the number of rate-limiting chemical transitions.

## 3 Materials and methods

### 3.1 Motor simulations

Kinetic Monte Carlo (KMC) methods, performed in Python, were used to model a Poisson stepper and obtain the motor trace *x*_*m*_(*t*), which was then used to obtain the bead trace *x*_b_(*t*) by integrating Eq (2). The input parameters were the mechanical step size *a*, the number *m* of (equally rate-limiting) chemical processes per mechanochemical step, the rate *k* for each of the *m* transitions, the stiffness *k*_*l*_ of the elastic linkage, and the radius *r*_*b*_ of the attached particle. The integration time was set to *t*_*c*_/15, where *t*_*c*_ = *γ/k*_*l*_ was the bead relaxation time.

### 3.2 Time domain analysis of position fluctuations

The displacement of the bead was calculated over integer numbers of points for each trace, and the variance of the displacement was plotted as a function of time. A weighted least square fit of Eq (7) was computed for a time interval of zero to 1/100 the trace duration, yielding the motor step size, *a*, and the linker stiffness, *k*_*l*_. Errors were calculated by performing 50 simulations with input parameters which matched the fit parameters of each of the experimental motors, then calculating the mean error of *a* and *k*_*l*_ over all the simulations.

### 3.3 Frequency domain analysis of velocity fluctuations

The normalized, one-sided, log-binned, power spectral density (PSD) of the bead’s speed was calculated for each trace. A least squares fit of Eq (13) to the PSD was performed to optimize the free parameters of motor step size, *a*, the stiffness of the system, *k*_*l*_, the number of rate-limiting steps, *m*, and for the experimental data, the drag of the system, *γ*. The average velocity of the motor, 〈*v*_m_〉, was calculated as the total angular distance traveled divided by the total time of the trace. Errors were calculated by performing 50 simulations with input parameters which matched the fit parameters of each of the experimental motors, then calculating the mean error of *a* and *k*_*l*_ over all the simulations.

## 4 Results

### 4.1 Simulated molecular motor rotation

Using simulations of a linear Poisson stepping motor (see Materials and Methods), we have tested the performance of each analysis method described above over a biologically relevant range of step sizes and linkage compliances. Fig 1 shows an example of such simulations and the analysis via the two methods. We performed fits of the simulated data, and the results of this analysis are shown in Fig 2. These simulations model a large load (1 *µ*m bead) attached to the motor; see SI, Fig S3, for similar simulations of a small load (30 nm bead) attached to the motor.

**Figure 1:**
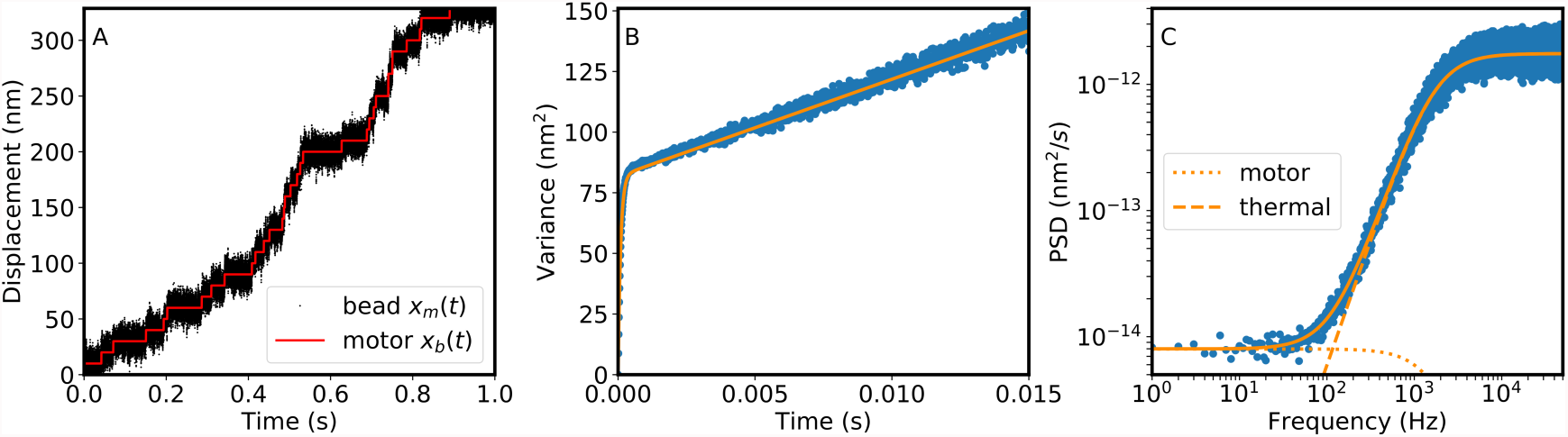
An example of the simulations used to test the three analysis methods. A) The simulated displacement versus time of a motor (red) and the attached bead (black). The first 1 s of a 60 s simulation is shown here; the full simulation was used for B-D. Simulation parameters: *r*_*b*_ = 500 nm, 〈*v*_m_〉 = 400 nm/s, *a* = 10 nm, *k*_*l*_ = 1 ⋅ 10^*−*5^ N/m. B) The variance of the reporter-particle displacement *var*(∆*x*_*b*_(*t*)) as a function of time for the simulated trace (blue points), and theory (Eq (7), orange line). C) The PSD of the speed of the bead (blue points) and theory (Eq (11), solid orange line). The dotted and dashed line show the components of the PSD attributed to the movement of the motor and the thermal noise on the bead, respectively. This PSD is an average of 60 PSDs taken on 1 s non-overlapping windows over the trace.

**Figure 2:**
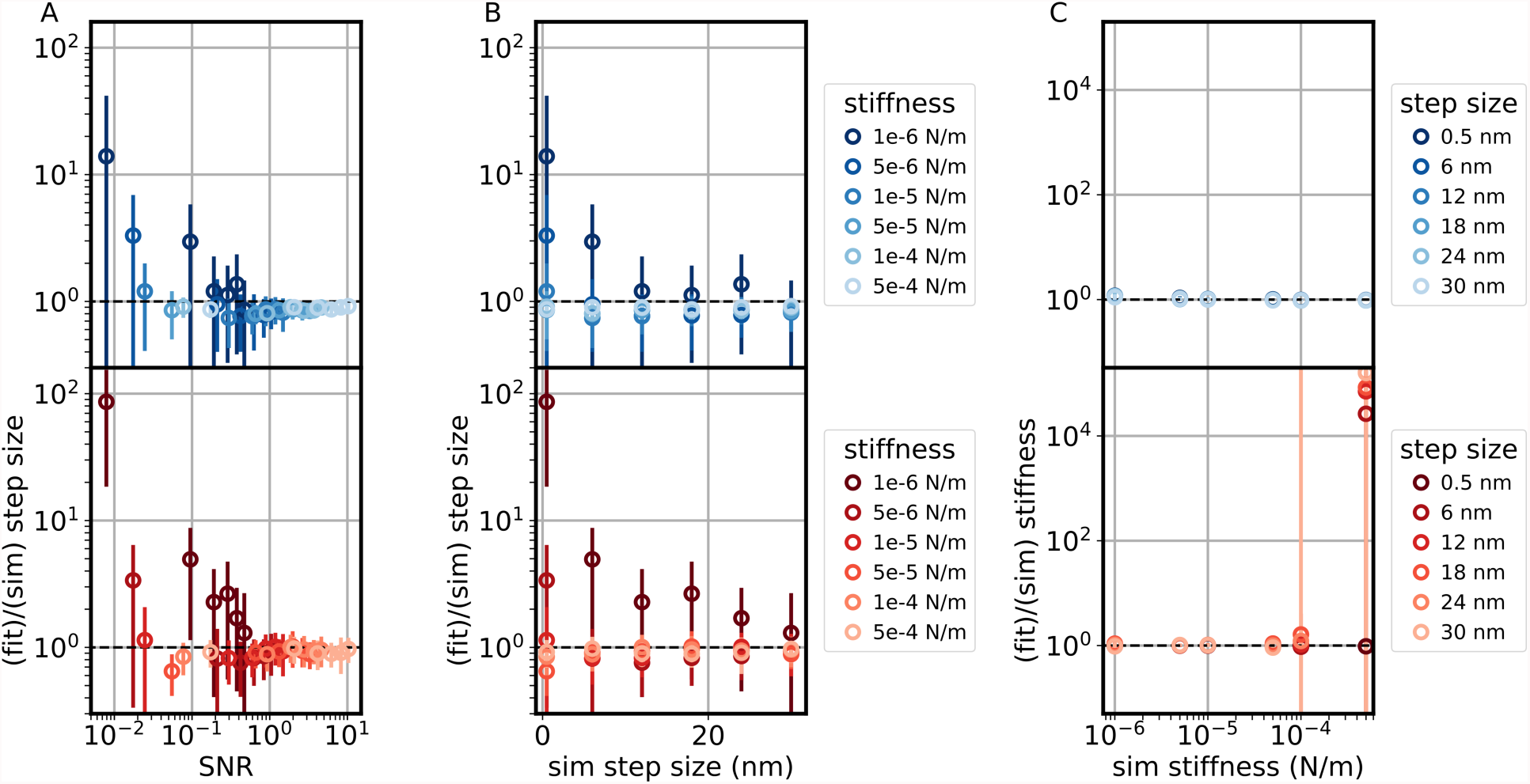
The performance of the frequency domain analysis of velocity fluctuations (blue) and time domain analysis of position fluctuations (red) analysis on simulated traces of a linear motor driving a 1 *µ*m bead at 400 nm/s with step size and stiffness as labeled. A) Ratio of the extracted step size to the simulated step size as a function of the signal to noise ration (SNR), defined as the simulated step size divided by the standard deviation of the difference between the bead position and motor position. B) Ratio of the extracted step size to the simulated step size as a function of the simulated step size. C) Ratio of the fit stiffness to the simulated stiffness as a function of the simulated stiffness. The dashed black lines represent perfect recovery of the input parameters. Points and error bars represent mean and standard deviation over 30 simulations (2 s each) per point.

We emphasize that the results shown in Fig 2 and Fig S3 are highly dependent upon the characteristics of both the motor and the experimental measurement. For example, with the frequency domain analysis of velocity fluctuations, recovery of the step size improves with longer measurements (yielding lower frequencies in the power spectrum), while recovery of the stiffness improves with higher acquisition rates (yielding higher frequencies in the power spectrum). Thus, simulations are a useful tool either to optimize such parameters prior to experiments or to explore under what circumstances these methods are applicable. In the following, we challenge the analysis methods developed above to extract the mechanical features of three molecular motors, whose parameters have been already independently assessed.

### 4.2 Kinesin

Kinesin is a cytoskeletal molecular motor which hydrolyzes ATP in order to move unidirectionally along microtubules, driving active cellular processes such as the transportation of molecular cargo. Moving with a ‘hand-over-hand’ technique, the heads of Kinesin-1 take 16 nm steps along the microtubule [25, 26]. Kinesin-1 is a model system for the study of molecular motors and development of single-molecule techniques [15, 27].

Fig 3A shows the translation of *Drosophila* kinesin-1 along an immobilized microtubule at saturating ATP concentration, tracked at 1 kHz via a gold nanoparticle (30 nm diameter) attached to one N-terminal head of the dimeric motor (data from Mickolajczyk et al [28]). Fig 3B shows the power spectrum (points) and fit (line) of the speed of the motor. The fit of the power spectrum yields a step size *a*=16 *±* 4 nm, where *m*=4 states are present per two mechanochemical cycles (as only one head is labeled), with a total stiffness *k*_*l*_=0.06 ± 0.05 pN/nm. The retrieved step is in agreement with the 16.4 nm step resolved in the time domain, and expected by the displacement of a single head along the microtubule. At saturating [ATP], the total duration of the mechanochemical cycle is evenly split between 1 head-bound (HB) and 2HB states [28], each of which contain one rate-limiting step [3, 29]. Thus, for saturating [ATP] assays in which a single head is labeled, one expects four rate-limiting processes per step, validating the value of *m* = 4 found here. Time domain analysis of position fluctuations (Fig S1B) yields a step size of 4.4 ± 0.9 nm; given four rate-limiting processes per step, this measurement is expected to be underestimated by a factor of 4, and is thus in agreement with the power spectral analysis. Finally, the total stiffness found from the fit in Fig 3B likely reflects a weighted average value of the stiffness of the system in the 1HB and 2HB states.

**Figure 3:**
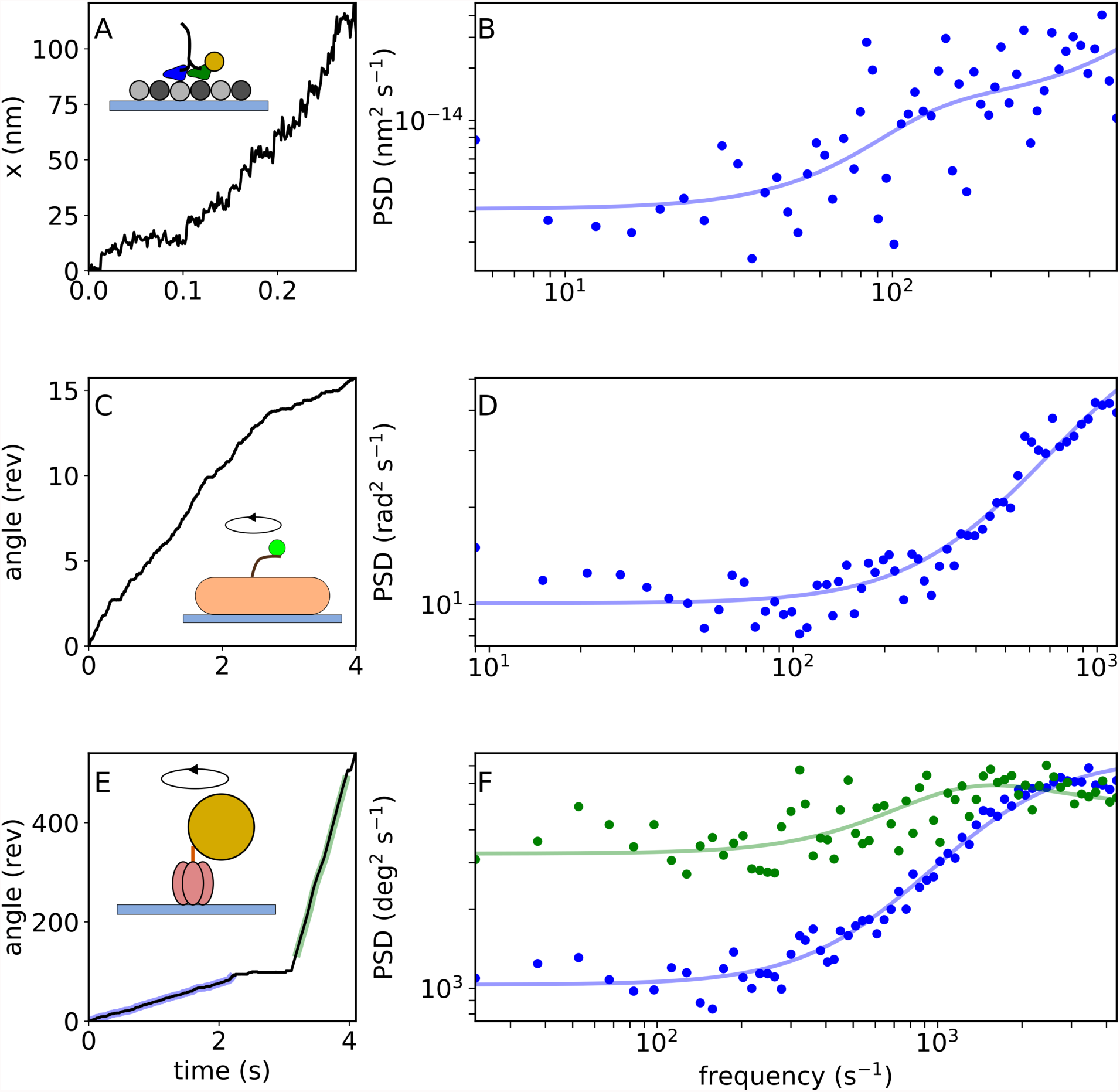
Experimental data of three molecular motors. A) Translation of *Drosophila* kinesin-1 along an immobilized microtubule, at saturating [ATP], tracked via a gold nanoparticle attached to one N-terminal head [27, 28]. Inset shows assay schematic. B) Points show the PSD of the speed of trace (A), and the line shows the best fit. (Fit parameters – *m* = 4; *a* = 16.3 nm; *k*_*l*_ = 0.06 pN /nm). C) Rotation of a bead attached to a truncated flagellar filament of a single BFM on a surface-immobilized *E. coli* cell [14]. Inset shows assay schematic. D) Points show the PSD of the speed of trace (C), and the line shows the best fit. (Fit parameters – *m* = 1; *a* = 11.7°; *k*_*l*_ = 1443 pN nm/rad). E) Rotation of a bead attached to the *γ* subunit of a single surface-immobilized yeast F_1_-ATPase [40]. Inset shows assay schematic. F) Points show the PSD of the speed of highlighted segments of trace (E), and the line shows the best fit. (Fit parameters – low [ATP] (blue): *m* = 1; *a* = 118°; *k*_*l*_ = 21 pN nm/rad; high [ATP] (green): *m* = 3; *a* = 95°; *k*_*l*_ = 23 pN nm/rad).

### 4.3 Bacterial flagellar motor

The bacterial flagellar motor (BFM) is the rotary molecular motor which rotates a flagellar filament of many species of motile bacteria, enabling swimming, swarming, and chemotaxis. Resolution of the fundamental step of the BFM has long proven difficult due to the rapid speed of motor rotation (up to 300 Hz in *E. coli* [30]), and the soft torsional compliance of the hook, which connects the rotor to the filament [31]. Using single molecule measurements of a particle attached to the filament of an immobilized bacterial cell, Sowa et al successfully resolved individual steps of the BFM under severely deenergized conditions [14].

Fig 3C shows the rotation of a polystyrene bead (227 nm diameter) attached to a truncated flagellar filament of a single BFM on a surface-immobilized *E. coli* cell (data from Sowa et al [14]). The speed of the motor was slowed via low expression of stator proteins and photodamage from the illumination source. Fig 3D shows the power spectrum of the speed of this trace (points) and theoretical best fit (line). The fit of the power spectrum yields a step size of 11.7 ± 0.5 degrees, equivalent to 31 ± 1 steps per revolution. This result is compatible with the results obtained by Sowa et al (13.8 degrees, 26 steps per revolution) [14], and also with the 26- or 34-fold symmetric ring of FliG molecules, the ‘track’ upon which the stators generate torque [32, 33]. Time domain analysis of position fluctuations (Fig S1C-D) yields a slightly larger step size, 38 ± 1 degrees, potentially affected by speed fluctuations during the trace (see SI). The fit of the power spectrum yields a stiffness of 1400 ± 400 pN nm/rad. Previous measurements have determined that the torsional compliance of the BFM is dominated by the compliance of the hook (instead of the filament) [31], and estimates of the torsional compliance of the hook span an order of magnitude, from 100 – 1300 pN nm rad^−1^ [31, 34, 35]. Our result is consistent with the upper range of these measurements.

### 4.4 F_1_ ATPase

F_1_F_*O*_ ATP synthase is the rotary molecular motor responsible for the synthesis of ATP in bacteria and eukaryotes. The enzyme is in fact comprised of two separable coaxial molecular motors, F_*O*_ and F_1_. On its own, F_1_ behaves as an ATP-hydrolyzing enzyme, the mechanochemical cycle of which has been well established by extensive single molecules studies [36]. Briefly, the rotation of F_1_ is comprised of six steps [11], of alternating size 80° and 40°. F_1_ consumes three ATP molecules per rotation (one for each of its catalytic sites); each ATP-binding dwell (three per rotation) shows single-exponential kinetics, and each catalytic dwell (also three per rotation) shows at least double-exponential kinetics, due to ATP hydrolysis and phosphate release [37–39].

Fig 3E shows the rotation of a 60 nm gold bead attached the the *γ* subunit of a single surface-immobilized yeast F_1_-ATPase (data from Steel et al [40]). The ATP concentration in the flow cell was changed from low to high (5 *µ*M to 3 mM) during the recording, evidenced by the change in rotation speed. Fig 3F shows the power spectra (points) and fit (line) of the highlighted sections of Fig 3E for low [ATP] and high [ATP] (blue and green, respectively). The fit of the low [ATP] spectrum yields 3.0 ± 0.4 steps per revolution, where *m* = 1 states are present, consistent with previous results which show 3 steps per revolution at low [ATP] [10], and show that this step of the mechanochemical cycle is limited by a single reaction, ATP-binding [11]. Time domain analysis of position fluctuations (Fig S1F-G) yields 2.7 ± 0.4 steps per revolution. The fit of Fig 3F for high [ATP] yields 3.8 ± 0.5 steps per revolution, where *m* = 3 states are present per mechanochemical cycle, yielding 11.4 ± 1.5 processes per revolution. Our model assumes that each step of the motor consists of the same number of Poisson processes, an assumption that doesn’t hold for F_1_ at non-limiting [ATP]. Nonetheless, the fit is consistent with 3-4 rate-limiting processes per catalytic site per revolution [41]. Time domain analysis of position fluctuations (Fig S1F-G) yields 7.2 ± 4 steps per revolution, also consistent with the number of rate-limiting mechanochemical processes. Interestingly, the fits of Fig 3F at low and high [ATP] yield a stiffness of 21 ±4 and 23 ± 3 pN nm/rad respectively, consistent with a previous measurement of 30 pN nm/rad and 66 pN nm/rad at the ATP-waiting and catalytic dwells, respectively [42].

## 5 Discussion

We have presented two new analytical solutions which are capable of extracting key features of the mechanochemical cycle of a molecular motor from single experimentally measured trajectories. We have used simulations to measure the performance of these methods under two quite different experimental conditions, and we have applied these methods to previously published data of three molecular motors.

Our simulations show that, with respect to recovering the step size, the stiffer the linker the better. This is apparent in both of the analysis methods, and most pronounced in the time domain analysis of position fluctuations, wherein a compliant linker lengthens the relaxation time of the system, introducing a correlation between dwell times. As stiffness increases, the methods perform similarly. Fig S3 compares these methods to a third method in which the distribution of the dwell time of the reporter particle over a defined window size allows the determination of the number of sequential steps within the window [43–46]. For our simulations, the two methods developed here recover the step size with better accuracy and lower error than the dwell time analysis method (see SI for discussion). We note that, while a stiffer linker improves recovery of step size, it may also increase the probability of interactions between the reporter-particle and motor which may disturb motor function.

Figs 2A and S3A show the recovery of the step size as a function of the ratio between the standard deviation of the noise and the step size. Crucially, these plots show good recovery of the step size up to and far beyond the limits of traditional step detection algorithms [47, 48], emphasizing the utility of these methods in situations where steps are not able to be resolved.

Simulations show that both the time domain analysis of position fluctuations and the frequency domain analysis of velocity fluctuations are able to recover the stiffness of the linker between the reporter particle and motor, showing little dependence on the step size. At both large and small loads, the two methods produce similar accuracy and error, at least for a compliant linker. As the stiffness increases, the accuracy of the time domain analysis of position fluctuations drops dramatically. The frequency domain analysis of velocity fluctuations performs better at higher stiffness, though simulations of a small load (where the corner frequency of the power spectrum shifts towards higher frequencies) show that recovery of the stiffness is affected by systematic errors associated with the discrete power spectra, such as aliasing and spectral leakage [49].

These two methods obviate the need for step-detection algorithms, which are often sensitive to user-set parameters [47, 50], and which prove of little use when the noise becomes of the order or larger than the step size. Furthermore, these methods are complementary, though we emphasize that simulations are useful to determine the accuracy and error of each method for a particular experiment. While both methods are based on the assumption of a Poisson stepper, where the time between two successive steps is exponentially distributed and the step size is constant, the frequency domain analysis of velocity fluctuations has the ability to further decipher the number of chemical processes per mechanical step in situations where the time between steps is a convolution of multiple exponentials. Both methods will be affected by off-pathway states, such as pauses, which are common amongst many molecular motors, though we note that this affects only the recovery of the step size, not the linker stiffness. Thus, the statistical analysis methods presented here provide a powerful mechanism of extracting kinetic features which are otherwise often invisible within single-molecule data.

## Acknowledgments

We thank KJ Mickolajczyk and WO Hancock for the kinesin data, Y Sowa and RM Berry for the BFM data, BC Steel and RM Berry for the F_1_ data, and all of the above for constructive discussion. FP and ALN acknowledge funding from the European Research Council under the European Unions Seventh Framework Programme (FP/2007-2013)/ERC Grant Agreement no. 306475. CBS is a member of the France-BioImaging (FBI) and the French Infrastructure for Integrated Structural Biology (FRISBI), 2 national infrastructures supported by the French National Research Agency (ANR-10-INBS-04-01 and ANR-10-INBS-05, respectively). MD

## Materials & Correspondence

To whom correspondence should be addressed

