## Supplementary Materials for "Kinetic analysis methods applied to single motor protein trajectories"

#### Derivations of equations

Below we provide derivations of various equations from the manuscript.

##### Variance of bead position: Eq (7)

Eq (2) from the manuscript can be rewritten as

$$dx_b(t) = \frac{k_l}{\gamma}(x_m(t) - x_b(t))\,dt + \frac{\xi(t)}{\gamma}\,dt, \tag{S1}$$

where

$$\begin{aligned} \langle \xi(t) \rangle &= 0 \\ \langle \xi(t)\xi(t') \rangle &= 2k_BT\gamma\,\delta(t-t') \\ t, t' > 0. & \end{aligned}$$

Rearranging (S1) and writing in an easily integratable form we have

$$d(x_b(t)e^{k_lt/\gamma}) = e^{k_lt/\gamma}\left(\frac{k_l}{\gamma}x_m(t) + \frac{\xi(t)}{\gamma}\right)dt\,.$$

Integrating from time 0 to time t gives

$$x_b(t) = x_b(0)e^{-k_lt/\gamma} + \int_0^t e^{-k_l(t-t')/\gamma}\left(\frac{k_l}{\gamma}x_m(t') + \frac{\xi(t')}{\gamma}\right)dt'\,. \tag{S2}$$

We are interested in calculating the stead-state variance,

$$\begin{aligned} var(\Delta t) &= \lim_{T\to\infty} \frac{1}{T}\int_0^T dt' \langle (x_b(t'+\Delta t) - x_b(t') - v\Delta t)^2\rangle \\ &= [\Delta x_b(t) = x_b(t) - v(t)] \\ &= \lim_{T\to\infty} \frac{1}{T}\int_0^T dt' \langle [\Delta x_b(t'+\Delta t) - x_b(t')]^2\rangle \\ &= \lim_{T\to\infty} \frac{1}{T}\int_0^T dt' [\langle \Delta x_b^2(t'+\Delta t)\rangle + \langle x_b^2(t')\rangle - 2\langle \Delta x_b(t'+\Delta t)\Delta x_b(t')\rangle]\rangle. \end{aligned}$$

We note that the integral will be dominated by large $t'$ as the integrand does not vanish as $t' \to \infty$, and thus,

$$var(\Delta t) = \langle \Delta x_b^2(\infty+\Delta t)\rangle + \langle x_b^2(\infty)\rangle - 2\langle \Delta x_b(\infty+\Delta t)\Delta x_b(\infty)\rangle.$$

To calculate this, it will be useful to know the cross-correlation,

$$\begin{aligned} \langle \Delta x_b(t)\Delta x_b(t')\rangle &= \langle (x_b(t) - vt)(x_b(t') - vt')\rangle \\ &= \langle x_b(t)x_b(t')\rangle - v^2tt', \end{aligned}$$

where

$$\langle x_b(t)x_b(t')\rangle = \\ \langle x_b^2(0)\rangle e^{-k_l(t+t')/\gamma} + \int_0^t dt_1 \int_0^{t'} dt_2\, e^{-k_l(t'-t_2)/\gamma - k_l(t-t_1)/\gamma} \Big\langle \Big(\frac{k_l}{\gamma}x_m(t_1) + \frac{\xi(t_1)}{\gamma}\Big)\Big(\frac{k_l}{\gamma}x_m(t_2) + \frac{\xi(t_2)}{\gamma}\Big)\Big\rangle,$$

where we have used the fact that neither motor position nor noise depend on the original position of the bead, $x_b(0)$, and that $\langle x_b(0)\rangle = 0$. To calculate the initial strength of the bead fluctuations we consider

$$dx_b(t) = -\frac{k_l}{\gamma}(x_b(t))\,dt + \frac{\xi(t)}{\gamma}\,dt, \langle \xi(t)\rangle = 0, \tag{S3}$$

where

$$\begin{aligned} \langle \xi(t)\rangle &= 0 \\ \langle \xi(t)\xi(t')\rangle &= 2k_BT\gamma\,\delta(t-t') \\ t, t' > 0, \end{aligned}$$

giving

$$\begin{aligned} \langle x_b^2(0)\rangle &= \int_{-\inf}^t dt' \int_{-\inf}^t dt''\, e^{-k_l(2t-t'-t'')/\gamma} \frac{\langle \xi(t')\xi(t'')}{\gamma^2} \\ &= \frac{2k_BT}{\gamma}\int_0^\infty dt'\, e^{-2k_lt'/\gamma} \\ &= \frac{k_BT}{k_l} \end{aligned}$$

If we assume that the motor position and the thermal noise are uncorrelated, $\langle \xi(t) x_m(t') \rangle = \langle \xi(t) \rangle \langle x_m(t') \rangle = 0$, then we have

$$\Big\langle \Big( \frac{k_l}{\gamma} x_m(t_1) + \frac{\xi(t_1)}{\gamma} \Big) \Big( \frac{k_l}{\gamma} x_m(t_2) + \frac{\xi(t_2)}{\gamma} \Big) \Big\rangle = \Big( \frac{k_l}{\gamma} \Big)^2 \langle x_m(t_1) x_m(t_2) \rangle + \frac{2 k_B T}{\gamma} \, \delta(t_1 - t_2) \, .$$

15 Applying Itô calculus to the motor position correlation and assuming $t_1 > t_2$,

$$\begin{aligned} \langle x_m(t_1) x_m(t_2) \rangle &= \int_0^{t_1} dt'_1 \int_0^{t_2} dt'_2 \langle \dot{x}_m(t'_1) \dot{x}_m(t'_2) \rangle \\ &= \sum_{t'_1} \Delta t \sum_{t'_2} \Delta t \langle \dot{x}_m(t'_1) \dot{x}_m(t'_2) \rangle \\ &= \sum_{t'_1 \neq t_2} \Delta t \Delta t \langle \dot{x}_m(t'_1) \rangle \langle \dot{x}_m(t'_2) \rangle + \sum_{t'_2} \Delta t \Delta t \langle \langle \dot{x}_m(t'_2) \rangle . \end{aligned}$$

16 As the terms in the first sum are regular as $t_1 \to t_2$, this becomes

$$\begin{aligned} \langle x_m(t_1) x_m(t_2) \rangle &= \int_0^{t_1} dt'_1 \int_0^{t_2} dt'_2 \langle \dot{x}_m(t'_1) \dot{x}_m(t'_2) \rangle + \sum_{t'_2} \Delta t^2 \langle \langle \dot{x}_m^2(t'_2) \rangle \\ &= v^2 t_1 t_2 + \sum_{t'_2} \Delta t^2 \langle \langle \dot{x}_m^2(t'_2) \rangle . \end{aligned}$$

17 The motor position is

$$x_m(t) = \sum_i a \delta(t - t_i), \tag{S4}$$

where $a$ is the size of the mechanical step of the motor, $t_i$ is the time at which the $i^{th}$ step is taken, distributed over $[0, \infty)$ with density $v/a$ (the number of steps per time). We then have

$$\sum_{t'_2} \Delta t^2 \langle \langle \dot{x}_m^2(t'_2) \rangle = \sum_{t'_2} \Delta t^2 \Big\langle \sum_{i,j} a^2 \, \delta(t_2 - t_i) \, \delta(t_2 - t_j) \Big\rangle .$$

18 This sum has zero values only when $i = j$, giving

$$\begin{aligned} \sum_{t'_2} \Delta t^2 \langle \langle \dot{x}_m^2(t'_2) \rangle &= \sum_{t'_2} \Delta t^2 \Big\langle \sum_i a^2 \, \delta(t_2 - t_i) \Big\rangle \\ &= \sum_{t'_2} \Delta t^2 \int_0^\infty dt_i \, \frac{v}{a} a^2 \delta^2(t_2 - t_i) \\ &= \sum_{t'_2} \Delta t^2 v a \, \delta(0) \\ &= \sum_{t'_2} \Delta t v a \\ &= v a t_2 \end{aligned}$$

In the above, we have used that when discrete $\delta(0) = 1/\Delta t$. Thus, the correlation of motor position is

$$\langle x_m(t_1)x_m(t_2)\rangle = v^2 t_1 t_2 + va\mathrm{min}(\mathrm{t_1}, \mathrm{t_2}).$$

Returning to the correlation of bead position, we have

$$\langle x_b(t)x_b(t'))\rangle = \frac{k_B T}{k_l} + A + B + C,$$

19 with

$$\begin{aligned} A &= \left(\frac{k_l v}{\gamma}\right)^2 \int_0^t dt_1\, e^{-k_l(t-t_1/\gamma)} t_1 \int_0^{t'} dt_2\, e^{-k_l(t'-t_2/\gamma)} t_2 \\ B &= \left(\frac{k_l}{\gamma}\right)^2 va \int_0^t dt_1 \int_0^{t'} dt_2\, e^{-k_l(t'-t_2)/\gamma - k_l(t'-t_1)/\gamma} \mathrm{min}(t_1, t_2) \\ C &= \int_0^t dt_1 \int_0^{t'} dt_2\, e^{-k_l(t'-t_2)/\gamma - k_l(t'-t_1)/\gamma} \frac{2k_B T}{\gamma}\, \delta(t_1 - t_2)\,. \end{aligned}$$

Considering the above term by term,

$$A = v^2 \Big[t - \frac{\gamma}{k_l}(1 - e^{-k_l t/\gamma})\Big] \Big[t' - \frac{\gamma}{k_l}(1 - e^{-k_l t'/\gamma})\Big].$$

20 Assuming $t > t'$, we can write

$$\begin{aligned} B &= \left(\frac{k_l}{\gamma}\right)^2 va \Big[\int_0^{t'} dt_2 \int_0^{t_2} dt_1 e^{-k_l(t'-t_2)/\gamma - k_l(t-t_1)/\gamma} t_1 + \int_0^{t'} dt_2 \int_{t_2}^{t} dt_1 e^{-k_l(t'-t_2)/\gamma - k_l(t-t_1)/\gamma} vat_2\Big] \\ &= av\Big(t' - \frac{\gamma}{2k_l}(1 - e^{-k_l t'/\gamma})^2 - \frac{\gamma}{2k_l}(1 - e^{-k_l t'/\gamma})(1 + e^{-k_l(t-t')/\gamma})\Big). \end{aligned}$$

21 Again assuming $t > t'$,the integral over $t_1$ in $C$ contributes nothing for $t_1 > t'$, as $t_2 < t'$ giving,

$$\begin{aligned} C &= \int_0^{t'} dt_1 \int_0^{t'} dt_2\, e^{-k_l(t'-t_2)/\gamma - k_l(t-t_1)/\gamma} \frac{2k_B T}{\gamma} \delta(t_1 - t_2) \\ &= e^{-k_l(t'+t)/\gamma} \frac{k_B T}{k_l} (e^{2k_l t'/\gamma} - 1) \\ &= \frac{k_B T}{k_l} (e^{-k_l(t-t')/\gamma} - e^{-k_l(t'+t)/\gamma}) \end{aligned}$$

22 Taken together, we have

$$\begin{aligned} \langle \Delta x_b(t+\Delta t)\Delta x_b(t)\rangle &= \langle x_b(t+\Delta t)x_b(t) - v^2 t(t+\Delta t) \\ &= \frac{k_B T}{k} - v^2(t+\Delta t)\frac{\gamma}{k_l}(1 - e^{-k_l t/\gamma}) - v^2 t \frac{\gamma}{k_l}(1 - e^{-k_l(t+\Delta t)/\gamma}) \\ &+ v^2 \Big[\frac{\gamma}{k_l}(1 - e^{-k_l(t+\Delta t)/\gamma})\Big] \Big[\frac{\gamma}{k_l}(1 - e^{-k_l/\gamma})\Big] \\ &- av\Big(t - \frac{\gamma}{2k_l}(1 - e^{-k_l t/\gamma})^2 - \frac{\gamma}{2k_l}(1 - e^{-k_l t/\gamma})(1 - e^{-k_l \Delta t/\gamma})\Big) \\ &+ \frac{k_B T}{k_l}(e^{-k_l \Delta t/\gamma} - e^{-k_l(2t+\Delta t)/\gamma}). \end{aligned}$$

As we are ultimately interested in the steady state regime, we will need

$$\langle \Delta x_b(\infty + \Delta t)\Delta x_b(\infty)\rangle =$$
$$\frac{k_BT}{k_l}(1 - e^{-k_l\Delta t/\gamma}) - v^2(\infty + \Delta t)\frac{\gamma}{k_l} - v^2\infty\frac{\gamma}{k_l} + \left(\frac{v\gamma}{k_l}\right)^2 - av(\infty + \Delta t - \frac{\gamma}{2k_l}(2 + e^{-k_l\Delta t/\gamma})),$$

as well as

$$\langle \Delta x_b^2(\infty + \Delta t)\rangle = \frac{2k_BT}{k_l} - 2v^2\frac{\gamma}{k_l}(\infty + \Delta t) + \left(\frac{v\gamma}{k_l}\right)^2 - av(\infty + \Delta t - \frac{3\gamma}{2k_l}),$$

and

$$\langle x_b^2(\infty)\rangle = \frac{2k_BT}{k_l} - 2v^2\frac{\gamma}{k_l}\infty + \left(\frac{v\gamma}{k_l}\right)^2 - av(\infty - \frac{3\gamma}{2k_l}).$$

Finally, bringing this all together we have,

$$\begin{aligned}
var(\Delta t) &= \langle \Delta x_b^2(\infty + \Delta t)\rangle + \langle x_b^2(\infty)\rangle - 2\langle \Delta x_b(\infty + \Delta t)\Delta x_b(\infty)\rangle \\
&= \frac{2k_BT}{k_l} - 2v^2\frac{\gamma}{k_l}(\infty + \Delta t) + (\frac{v\gamma}{k_l})^2 - av(\infty + \Delta t - \frac{3\gamma}{2k_l}) + \frac{2k_BT}{k_l} - 2v^2\frac{\gamma}{k_l}\infty \\
&\quad + \left(\frac{v\gamma}{k_l}\right)^2 - av(\infty - \frac{3\gamma}{2k_l}) - \frac{2k_BT}{k_l}(1 - e^{-k_l\Delta t/\gamma}) + 2v^2\frac{\gamma}{k_l}(\infty + \Delta t) + 2v^2\frac{\gamma}{k_l}\infty \\
&\quad - (2\frac{v\gamma}{k_l})^2 + 2av(\infty + \Delta t - \frac{\gamma}{2k_l}(2 + e^{-k_l\Delta t/\gamma})) \\
&= \frac{2k_BT + av\gamma}{k_l}(1 - e^{-k_l\Delta t/\gamma}) \qquad \text{(S5)}
\end{aligned}$$

#### **PSD of the speed of the bead: Eq** (8)

The PSD of the function $f(t)$ is defined by

$$PSD_f(\omega) = \lim_{T\to\infty}\frac{1}{T} < |\mathcal{F}[f](\omega)|^2 > \qquad \text{(S6)}$$

where the Fourier transform of $f(t)$ is

$$\mathcal{F}[f](\omega) = \int_{-T/2}^{T/2} f(t)e^{-i\omega t}. \qquad \text{(S7)}$$

The equation of motion for the bead position $x_b$, given the position of the motor $x_m$ and the thermal noise $\xi(t)$ is (Eq. (2)):

$$\gamma\dot{x}_b(t) = kx_m(t) - kx_b(t) + \xi(t) \qquad \text{(S8)}$$

The Fourier transform of Eq. (S8) is

$$i\omega\gamma\mathcal{F}[x_b](\omega) = k\mathcal{F}[x_m] - k\mathcal{F}[x_b] + \mathcal{F}[\xi] \qquad \text{(S9)}$$

So

$$\mathcal{F}[x_b](\omega) = \frac{k\mathcal{F}[x_m] + \mathcal{F}[\xi]}{k + i\omega\gamma}$$

Reminder:

$$\mathcal{F}[\frac{df}{dt}] = i\omega\mathcal{F}[f]$$

given by integration by parts and by

$$[f(t)e^{-i\omega t}]_{-\infty}^{+\infty} = 0$$

The PSD of the velocity of the bead $v_b = dx_b/dt$ is then (where $v_m = dx_m/dt$ is the velocity of the motor)

$$PSD_{v_b}(\omega) \tag{S10}$$

$$= \lim_{T\to\infty} \frac{1}{T} < |\mathcal{F}[\frac{dx_b}{dt}](\omega)|^2 > \tag{S11}$$

$$= \lim_{T\to\infty} \frac{1}{T} < |i\omega\mathcal{F}[x_b]|^2 > \tag{S12}$$

$$= \lim_{T\to\infty} \frac{1}{T} < \left| i\omega \frac{k\mathcal{F}[x_m] + \mathcal{F}[\xi]}{k + i\omega\gamma} \right|^2 > \tag{S13}$$

$$= \lim_{T\to\infty} \frac{1}{T} < \left| \frac{k\mathcal{F}[\frac{dx_m}{dt}] + i\omega\mathcal{F}[\xi]}{k + i\omega\gamma} \right|^2 > \tag{S14}$$

$$= \lim_{T\to\infty} \frac{1}{T} < \left| \frac{k\mathcal{F}[\frac{dx_m}{dt}]}{k + i\omega\gamma} + i\omega \frac{\mathcal{F}[\xi]}{k + i\omega\gamma} \right|^2 > \tag{S15}$$

$$= \lim_{T\to\infty} \frac{1}{T} (< \left| \frac{k\mathcal{F}[\frac{dx_m}{dt}]}{k + i\omega\gamma} \right|^2 > + < \left| i\omega \frac{\mathcal{F}[\xi]}{k + i\omega\gamma} \right|^2 > + \tag{S16}$$

$$+2 < \frac{k\mathcal{F}[\frac{dx_m}{dt}]}{k + i\omega\gamma} \frac{i\omega\mathcal{F}[\xi]}{k + i\omega\gamma} > \tag{S17}$$

$$= \frac{k^2}{k^2 + \omega^2\gamma^2} PSD_{v_m}(\omega) + \frac{\omega^2}{k^2 + \omega^2\gamma^2} PSD_{\xi}(\omega) \tag{S18}$$

#### **PSD of motor velocity for more than one rate-limiting step: Eq** (12)

The power spectrum of a signal $v(t)$ can be written in terms of the two-time correlation function (Wiener-Khinchin theorem)

$$R_v(\tau) = \langle v(t)v(t-\tau) \rangle$$

$$S_v(\omega) = R_v(\omega) = \int_{-\infty}^{\infty} e^{i\omega t} R_v(\tau) dt$$

For a discrete stepping motor, $v(t) = \sum_{-\infty}^{\infty} d\delta(t - t_n)$, we can write

$$
\begin{aligned}
R_v(\omega) &= lim_{T\to\infty}\frac{d^2}{T}\sum_{n,m=-\infty}^{\infty}\int_{-\infty}^{\infty} e^{i\omega t}R_v(\tau)\langle\delta(t_n - t_m - \tau)\rangle dt \\
&= lim_{T\to\infty}\frac{d^2}{T}\sum_{n,\Delta n=-\infty}^{\infty}\int_{-\infty}^{\infty} e^{i\omega t}R_v(\tau)\langle\delta(\Delta t_{\Delta n} - \tau)\rangle dt \\
&= lim_{T\to\infty}\frac{d^2}{T}\frac{T}{\langle\Delta t_1\rangle}\sum_{\Delta n=-\infty}^{\infty}\int_{-\infty}^{\infty} e^{i\omega t}R_v(\tau)\langle\delta(\Delta t_{\Delta n} - \tau)\rangle dt \\
&= \frac{d^2}{\langle\Delta t_1\rangle}\sum_{\Delta n=-\infty}^{\infty}\int_{-\infty}^{\infty} e^{i\omega t}R_v(\tau)\langle\delta(\Delta t_{\Delta n} - \tau)\rangle dt \\
&= \frac{d^2}{\langle\Delta t_1\rangle}\sum_{\Delta n=-\infty}^{\infty}\langle e^{i\omega\Delta t_{\Delta n}}\rangle \\
&= \frac{d^2}{\langle\Delta t_1\rangle}(1 + \sum_{\Delta n=1}^{\infty}(\langle e^{i\omega\Delta t_{\Delta n}}\rangle + \langle e^{-i\omega\Delta t_{\Delta n}}\rangle)) \\
&= \frac{d^2}{\langle\Delta t_1\rangle}(1 + Q(\omega) + Q(-\omega)
\end{aligned}
$$

where $\Delta n = m - n$, $\Delta t_{\Delta n} = t_n - t_m$, and $\frac{T}{\langle\Delta t_1\rangle}$ is the number of steps in time $T$, $Q(\omega) = \sum_{\Delta n=1}^{\infty} q_{\Delta n}(\omega)$, and $q_{\Delta n}(\omega) = \langle e^{i\omega\Delta t_{\Delta n}}\rangle$. This general formula applies to any stepping motor characterized by a $Q(\omega)$ according to

$$
\begin{aligned}
q_{\Delta n}(\omega) &= \langle e^{i\omega\Delta t_{\Delta n}}\rangle \\
&= \int_0^{\infty} P_{\Delta n}(\Delta t)e^{i\omega\Delta t}d\Delta t
\end{aligned}
$$

If the motor is a Poisson stepper, the probability that the motor takes $N$ steps in time $t$, each step of average duration $\tau_0$, is given by the Poisson distribution,

$$P_{\Delta n}(\Delta t) = \frac{(\Delta t^{\Delta n-1}}{(\Delta n - 1)!\Delta t_1^{\Delta}n}e^{-(\Delta t/\Delta t_1}, \tag{S19}$$

and

$$Q(\omega) + Q(-\omega) + 1 = 1$$

If instead we have a stepping motor that takes $m$ exponential sub-steps of average duration $\tau_0/m$, the probability is now,

$$P_{\Delta n,m}(\Delta t) = \frac{(\Delta t^{\Delta n(m-1)}}{\Delta n(m-1)!(\Delta t_1/m)^{\Delta}n}e^{-(\Delta t/(\Delta t_1/m)} \tag{S20}$$

and

$$
\begin{aligned}
q_{\Delta n}(\omega) &= \langle e^{i\omega t_N} \rangle \\
&= \int_0^\infty P_{\Delta n,m}(\Delta t) e^{i\omega \Delta t} d\Delta t \\
&= \int_0^\infty \frac{\Delta t^{\Delta n(m-1)}}{\Delta n(m-1)!(\tau_0/m)^{\Delta n}} e^{-\Delta t/(\tau_0/m)} e^{i\omega\Delta t} d\Delta t \\
&= \frac{1}{(1 - iw\langle \Delta t_1 \rangle/m)^{\Delta nm}} \\
Q(\omega) &= \sum_{N=1}^{\infty} q_N(\omega) \\
&= \sum_{N=1}^{\infty} \frac{1}{(1 - iw\langle \Delta t_1 \rangle/m)^{Nm}} \\
&= \frac{1}{(1 - iw\langle \Delta t_1 \rangle/m)^{m} - 1} \\
Q(\omega) + Q(-\omega) + 1 &= \frac{1}{(1 - iw\langle \Delta t_1 \rangle/m)^{m} - 1} + \frac{1}{(1 + iw\langle \Delta t_1 \rangle/m)^{m} - 1}
\end{aligned}
$$

And thus,

$$
\begin{aligned}
S_{v_m} &= \frac{d^2}{\langle \Delta t_1 \rangle} \alpha \\
&= d\langle v_m \rangle \alpha
\end{aligned}
$$

where

$$
\alpha = \left( 1 + \frac{1}{(1 - 2\pi i f \tau_1/m)^m - 1} + \frac{1}{(1 + 2\pi i f \tau_1/m)^m - 1} \right). \tag{S21}
$$

### Comparison to time domain dwell time analysis

For the process where $m$ "hidden" sequential chemical steps are required to produce one observable mechanical step, if the transition between each internal state is described by a single rate constant, $k_i = k$, the dwell time distribution, as calculated by convolution of $m$ Poissonian processes [1], is described by the Gamma distribution (or more precisely, given that $m > 0$ is an integer, the Erlang distribution)

$$
p(\tau) = \frac{k^m \tau^{m-1}}{(m-1)!} e^{-k\tau}. \tag{S22}
$$

Thus, fitting experimentally measured dwell time distributions to Eq (S22) allows the determination of the number of sequential steps, assuming the rate constants for each step are comparable [1–4]. In a scenario where the signal to noise allows resolution of individual mechanical steps, this analysis provides the number of rate-limiting biochemical processes per mechanical step. For experiments in which individual mechanical steps are not resolved, this analysis may be applied on an arbitrary distance $\Delta x$ over which the dwell time can be reliably measured, thereby yielding the number of rate-limiting biochemical and mechanical steps over this distance [5, 6]. In case the experimental dwell time distribution differs from the one provided by Eq (S22), its shape can be used to suggest and build a more complete kinetic model for the motor [7, 8]. On the other hand, this treatment does not provide a way to measure or estimate the stiffness of the system.

This analysis method was applied to the simulations described in Materials and Methods. A dwell time window was defined as three times the estimated step size. For each trace, dwell times were calculated as the time to traverse the dwell time window. A normalized histogram of dwell times was fit with a gamma function, as per Eq (S22). The step size of the motor $a$ was then calculated from the rate constant from the fit. Errors were calculated by performing 50 simulations with input parameters which matched the fit parameters of each of the experimental motors, then calculating the mean error of $a$ over all the simulations.

We emphasize that the results shown in Fig 2 and Fig S3 are highly dependent upon the characteristics of both the motor and the experimental measurement. The time domain dwell time analysis shows improved recovery of the step size for larger window sizes, given an infinitely long trace. For finite traces, a larger window size yields fewer points to fit Eq (S22), so there exists an optimal window size. For the parameters used in these simulations, both the time domain analysis of positions flucturations and the frequency domain analysis of speed fluctuations show better accuracy and precision in the recovery of the motor step size.

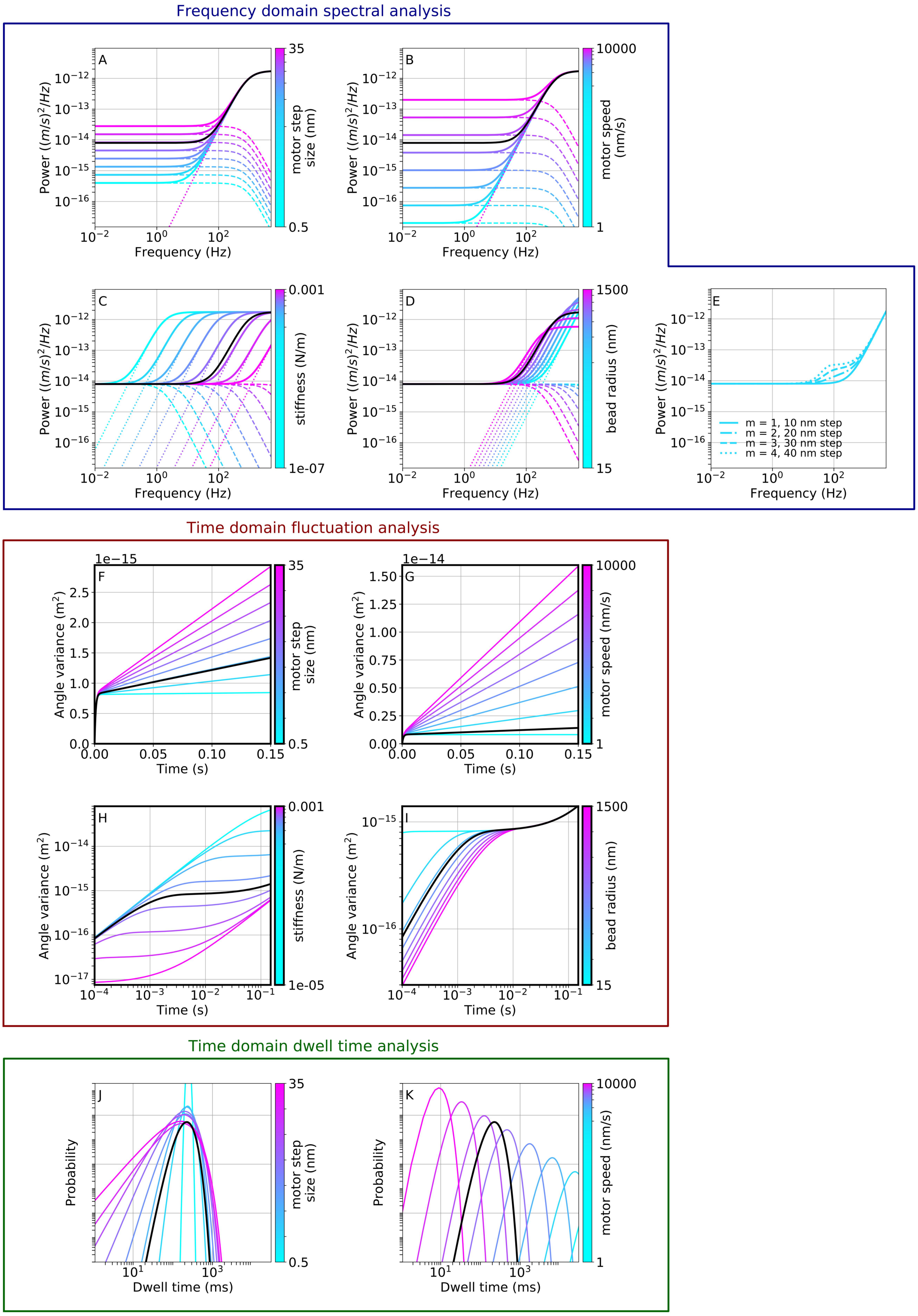


Figure S1: Demonstration of the theoretical results from the three analysis methods. A-E) Frequency domain spectral analysis, F-I) Time domain fluctuation analysis, J-K) Time domain dwell time analysis. For comparison, the black line in each subplot represents a 1 $\mu$m bead translocated at a speed of 400 nm/s by a motor which takes 10 nm steps, and a torsional hook stiffness of $1{\cdot}10^{-5}$ N/m. Each subplot varies a single variable from this standard, either linear or log-spaced, as shown and labeled by the colorbar on the right of the legend. Plot (E) shows the affects of the number of kinetic states for a 60 nm bead.

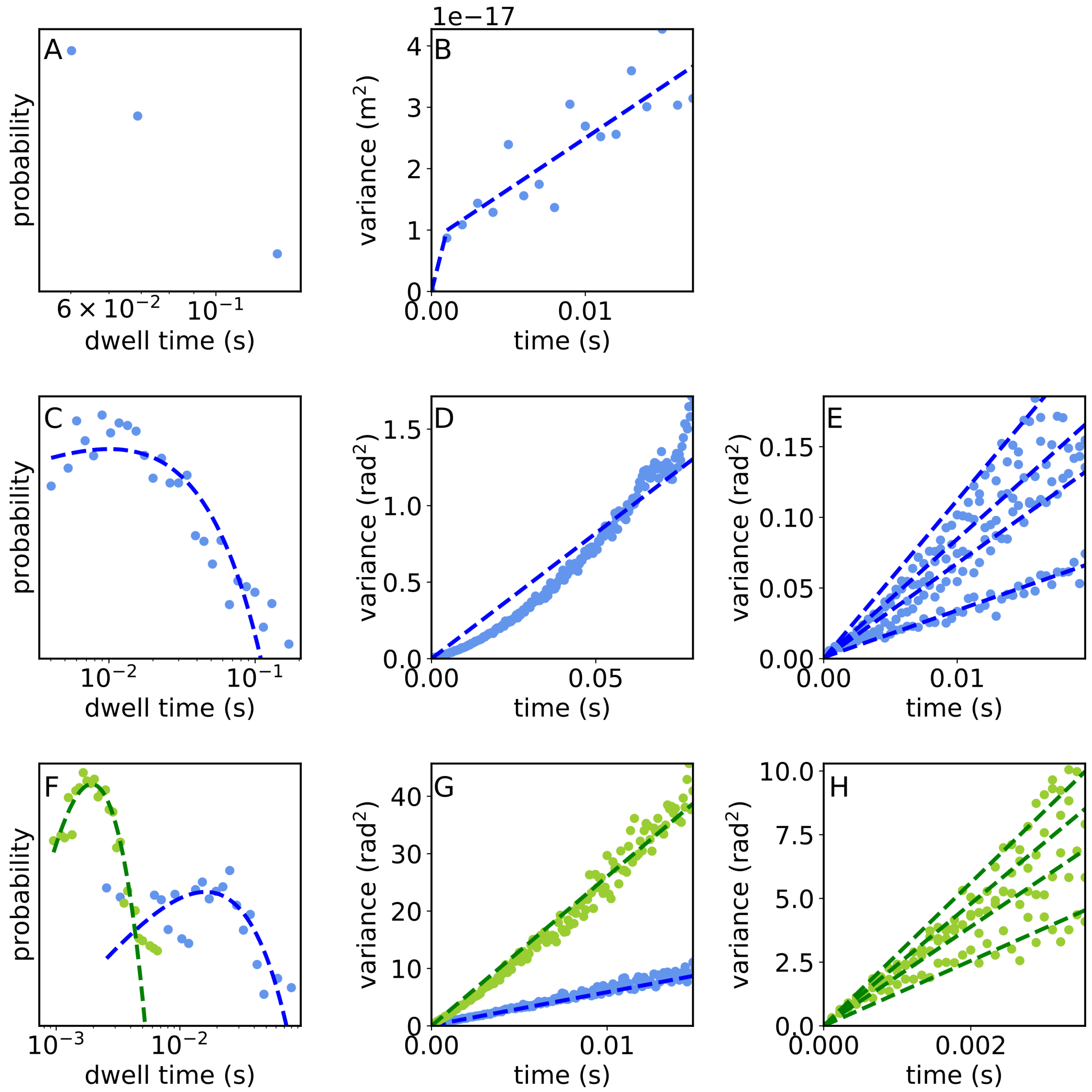


Figure S2: Experimental data (points) and fits (dashed lines) of the three molecular motors presented in Fig 3 of the manuscript, analyzed with the time domain dwell time analysis and time domain analysis of position fluctuations. A) The length of the kinesin trace does not enable a fit to the dwell time data. B) time domain analysis of position fluctuations of kinesin yields a step size of 4 $\pm$ 1 nm and a stiffness of (1.0 $\pm$ 0.3 )$\cdot 10^{-3}$ N/m. C) The dwell time analysis of the BFM yields a step size of 22 $\pm$ 3 degrees D) The time domain analysis of position fluctuations of the BFM yields a step size of 38 $\pm$ 2 degrees and a stiffness larger than can be reliably fit (greater than 300 pN nm/rad for the experimental conditions, as determined via simulation, not shown). E) The same analysis as in (D), but performed on four non-overlapping subsections of the trace, yielding step sizes of 15-18 degrees. The concavity in (D) arises from changes in motor speed. F) Dwell time analysis of $F_1$ yields a step size of 127 $\pm$ 11 degrees and 35 $\pm$ 8 degrees at low [ATP] (blue) and high [ATP] (green), respectively. G) Time domain analysis of position fluctuations of $F_1$ yields a step size of 132 $\pm$ 14 degrees and 50 $\pm$ 12 degrees at low [ATP] (blue) and high [ATP] (green), respectively. The stiffness is too high to be reliably fit (greater than 1 pN nm/rad for the experimental conditions, as determined via simulation, not shown). H) The same analysis as in (G), but performed on four non-overlapping subsections of the trace, yielding step sizes of 116-220 and 25-55 at low and high [ATP].

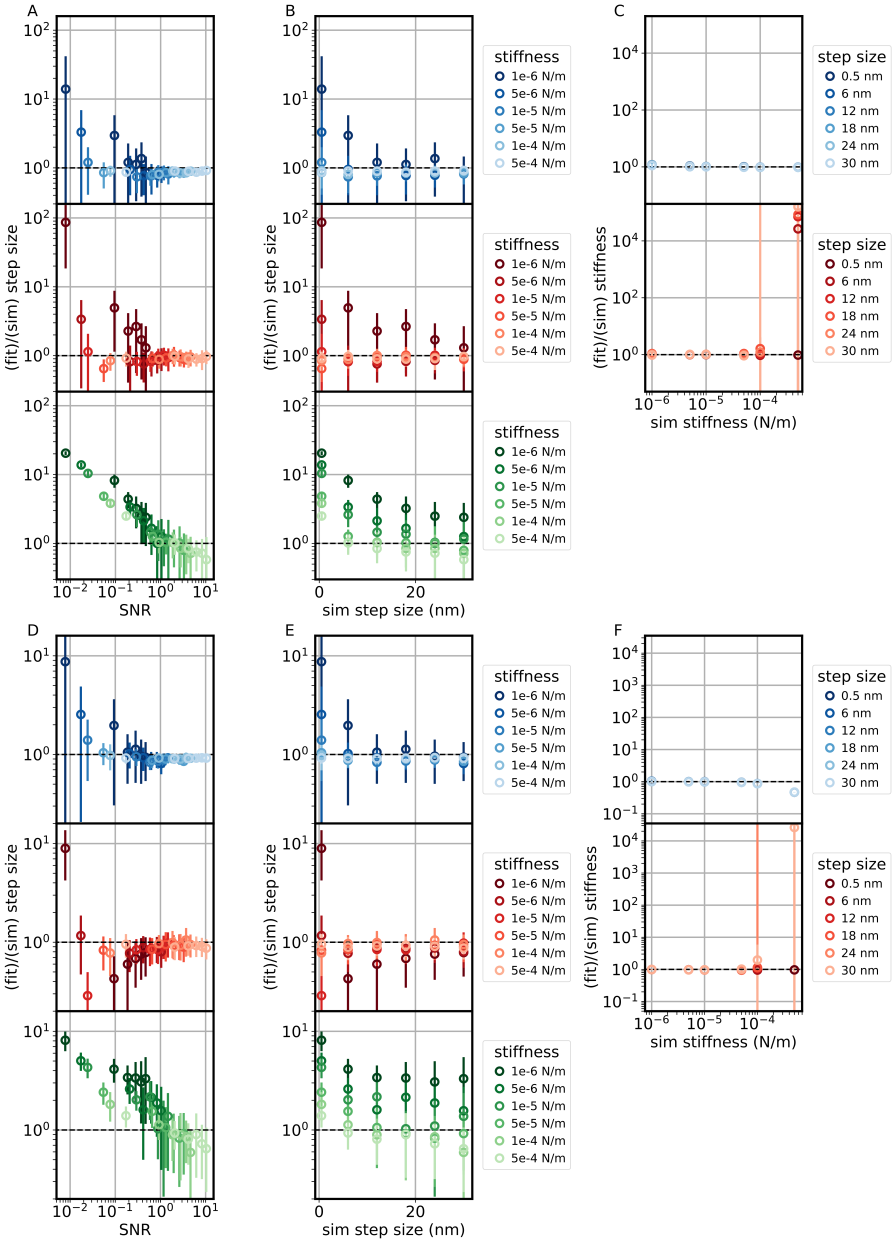


Figure S3: The performance of the frequency domain analysis of velocity fluctuations (blue) and time domain analysis of position fluctuations (red), compared to the time domain analysis of dwell times (green, Eq (S22)) on simulated traces of a linear motor driving A-C) a 1 $\mu$m bead at 400 nm/s or D-F) a 30 nm bead at 400 nm/s with step size and stiffness as labeled. A,D) Ratio of the extracted step size to the simulated step size as a function of the signal to noise ration (SNR), defined as the simulated step size divided by the standard deviation of the difference between the bead position and motor position. B,E) Ratio of the extracted step size to the simulated step size as a function of the simulated step size. C,F) Ratio of the fit stiffness to the simulated stiffness as a function of the simulated stiffness. The dashed black lines represent perfect recovery of the input parameters. Points and error bars represent mean and standard deviation over 30 simulations (2 s each) per point.
